## supplemental Table and Figures for "C-type natriuretic peptide improves maternally aged oocytes quality by inhibiting excessive PINK1/Parkin-mediated mitophagy": Supplemental Materials.docx

**Running title:** CNP improves aged oocyte quality

**Hui Zhang^1, 2a^, Chan Li^1, 2a^, Qingyang Liu^1, 2^, Jingmei Li^1, 2^, Hao Wu^1, 2^, Rui Xu^1, 2^, Yidan Sun^1, 2^, Ming Cheng^1, 2^, Xiaoe Zhao^1, 2^, Menghao Pan^1, 2^, Qiang Wei^1, 2b^, Baohua Ma^1, 2b^**

1 College of Veterinary Medicine, Northwest A&F University, Yangling, Shaanxi, People’s Republic of China,

2 Key Laboratory of Animal Biotechnology, Ministry of Agriculture, Yangling, Shaanxi, People’s Republic of China

**^a^** These authors contributed equally.

**^b^** Corresponding author: Baohua Ma and Qiang Wei

**Address:** College of Veterinary Medicine, Northwest A&F University, Yangling, Shaanxi, People’s Republic of China; Key Laboratory of Animal Biotechnology, Ministry of Agriculture, Yangling, Shaanxi, People’s Republic of China. 712100

**Table S1 Primers sequences**

| Gene name | Primer sequences | Amplicon size (bp) | NCBI Reference Sequence | |
| --- | --- | --- | --- | --- |
| *Gapdh* | F: 5’-TCACTGCCACCCAGAAGA-3’  R: 5’-GACGGACACATTGGGGGTAG-3’ | 185 | | XM_017321385.2 |
| *Zfp640* | F: 5’-TGTGCAGGCTTGAATGGTTC-3’  R: 5’-CTGAGGTCCCCTCAATGCAC-3’ | 70 | | XM_030247511.2 |
| *Gm6749* | F: 5’-ATGGGTGTTGCCCATACCAC-3’  R: 5’-GGCTTTTGTGGCCAGTTGTT-3’ | 93 | | XM_017321875.3 |
| *Obox7* | F: 5’-TTGTCCGCAAGAATACCAAGAA-3’  R: 5’-AGAGCTTGTCTGCAGATGGAC-3’ | 188 | | NM_001038676.1 |
| *Glrx* | F: 5’-CATAGGCGGATGCAGTGATCT-3’  R: 5’-CTCTGCCTGCCACCCCTTTTAT-3’ | 106 | | NM_053108.4 |
| *Vamp9* | F: 5’-ACTCTCTTTATTGACGGAATCACT-3’  R: 5’-TATTCCATCATGTGCTGTGCT-3’ | 194 | | NM_001378420.1 |
| *Tnk2* | F: 5’-CAAGAGGGCCAGGTAGTGTG-3’  R: 5’-TTGGCACTAGAGCAACCCTG-3’ | 170 | | NM_016788.3 |
| *Cd72* | F: 5’-CCTCGGAAGTCTGGAGGAGA-3’  R: 5’-GGGGCGTCAGAGAGGTATTC-3’ | 108 | | NM_001110320.1 |

**Figure S1**

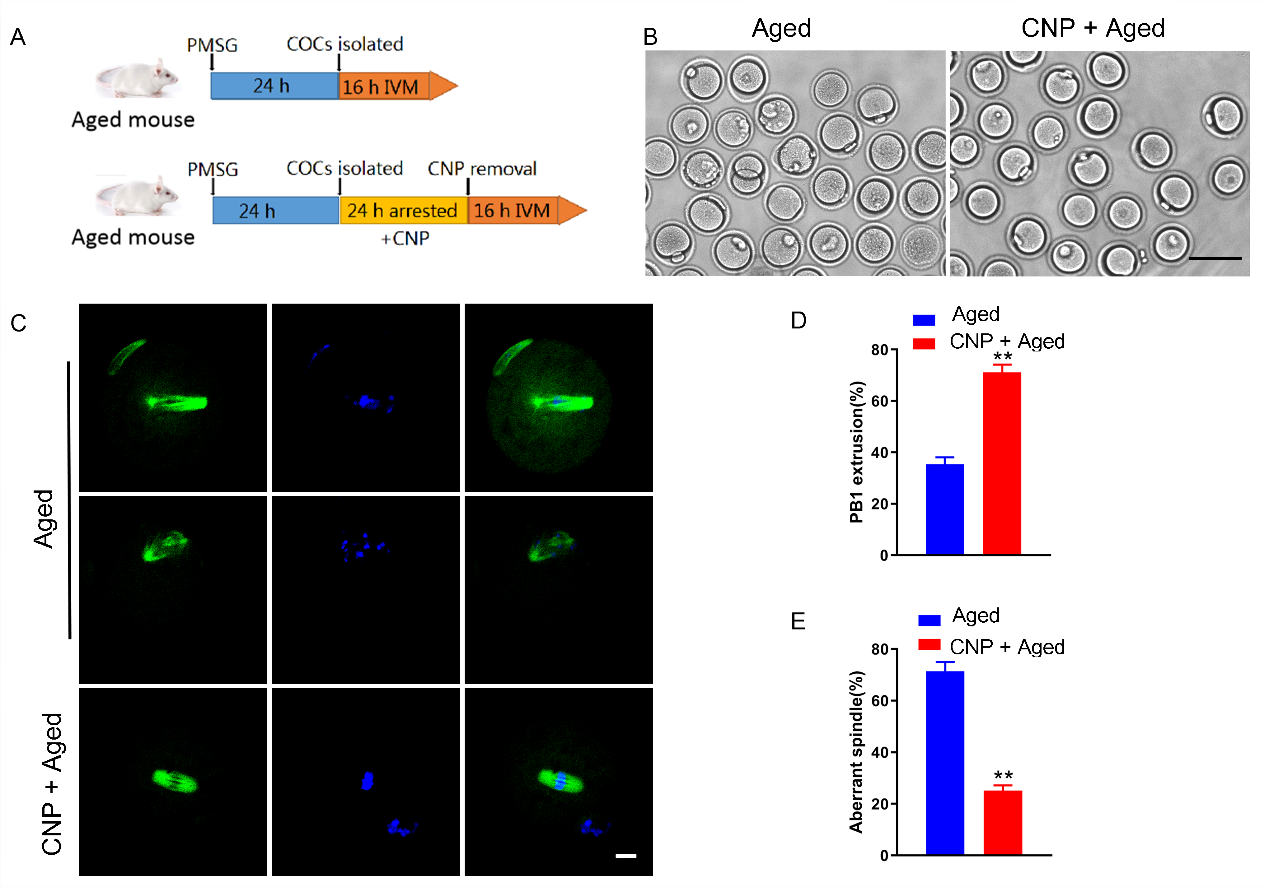

**Figure S1. Effects of CNP on the maturation and spindle/chromosome structure in aged oocytes.** (A) A timeline diagram of experimental design. (B) Representative images of the oocyte polar body extrusion in aged control and CNP treatment groups. Scale bar, 100 μm. (C) Rate of polar body extrusion in aged control and CNP treatment groups. **, significant difference (P < 0.01). (D) Representative images of the spindle morphology and chromosome alignment at metaphase II in aged control and CNP treatment oocytes. Scale bar, 10 μm. (E) The rate of aberrant spindles at metaphase II was recorded in aged control and CNP treatment oocytes. **, significant difference (P < 0.01)

**Figure S2**

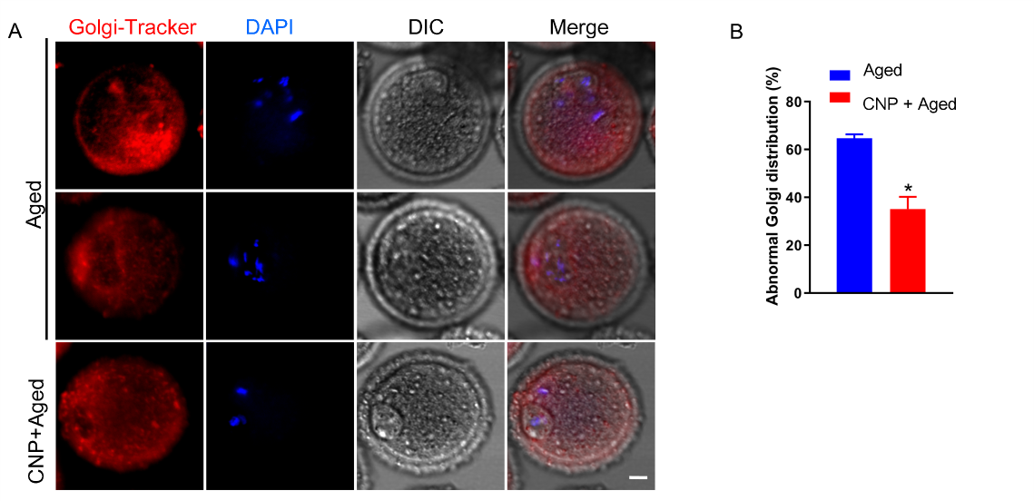

**Figure S2. Effects of CNP on Golgi apparatus distribution in aged oocytes.** Representative images of Golgi apparatus distribution in the aged control and CNP treatment oocytes stained with Golgi-Tracker Red. Scale bar, 10 μm. (B) Abnormal distribution of Golgi apparatus in the aged control and CNP treatment oocytes. *, significant difference (P< 0.05).

**Figure S3**

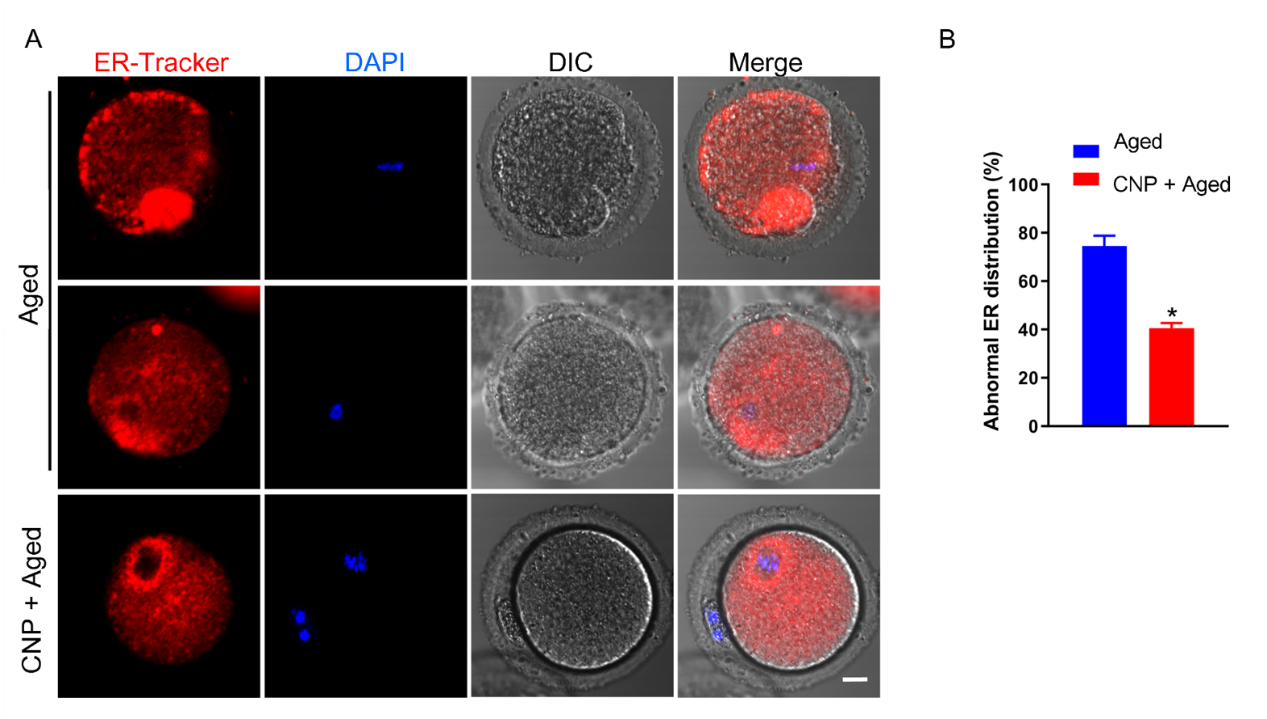

**Figure S3. Effects of CNP on endoplasmic reticulum distribution and function in aged oocytes.** (A) Representative images of ER distribution in the aged control and CNP treatment oocytes stained with ER-Tracker Red. Scale bar, 10 μm. (B) Abnormal distribution of ER in the aged control and CNP treatment oocytes. *, significant difference (P < 0.05).

**Figure S4**

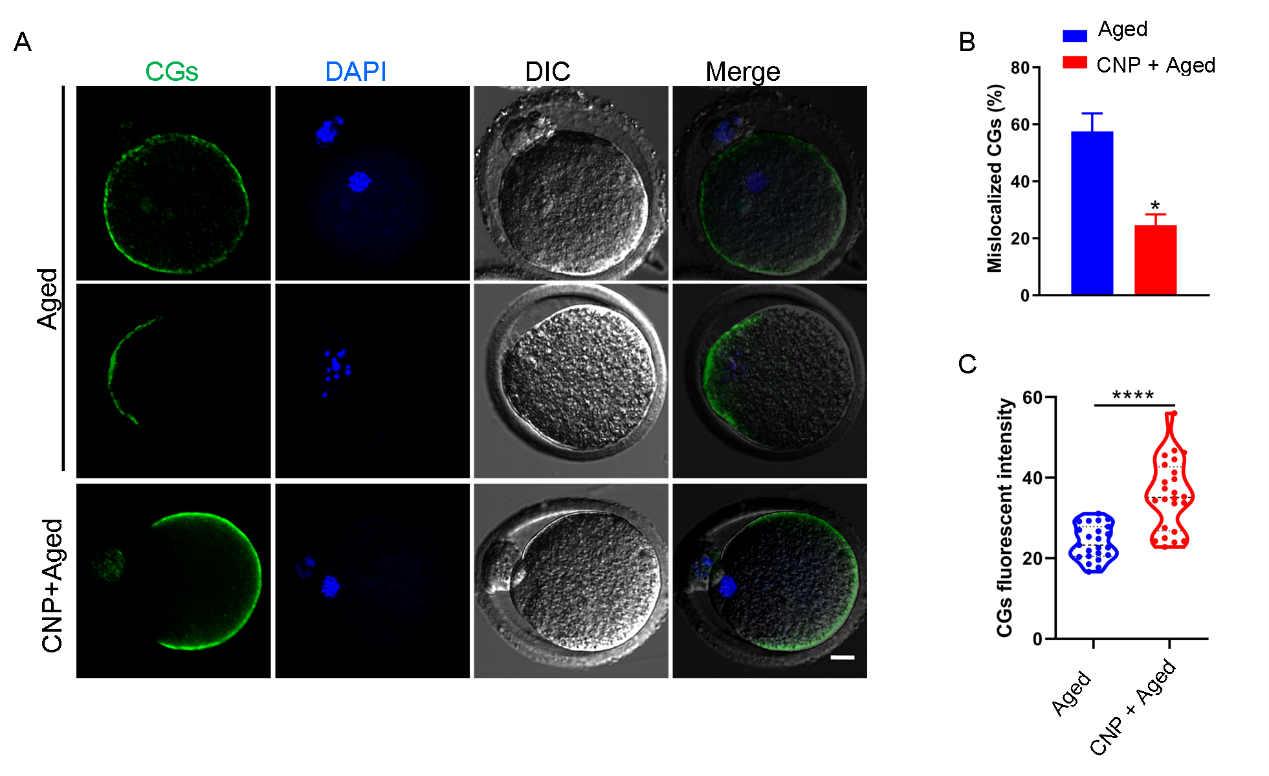

**Figure S4. Effects of CNP on the dynamics of CGs in aged oocytes.** Representative images of CGs distribution in the aged control and CNP treatment oocytes stained with LCA-FITC and imaged by confocal microscope. Scale bar, 10 μm. (B) The rate of mislocalized CGs was recorded in the aged control and CNP treatment oocytes. *, significant difference (P < 0.05) (C) The fluorescence intensity of CG signals was measured in the aged control and CNP treatment oocytes. ****, significant difference (P < 0.0001)

**Figure S5**

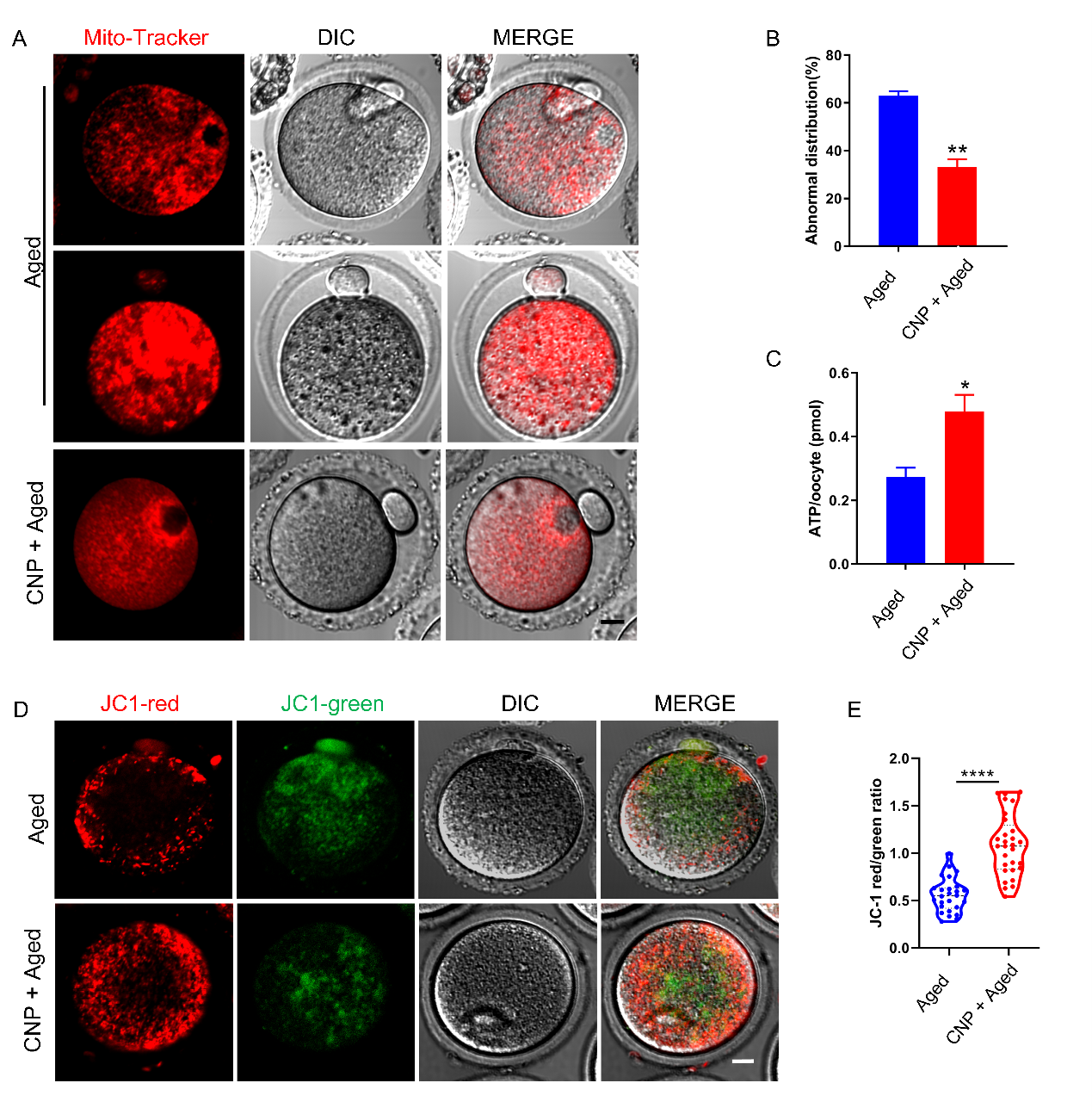

**Figure S5. Effects of CNP on the mitochondrial distribution and function in aged oocytes.** (A) Representative images of mitochondrial distribution in aged control and CNP treatment oocytes stained with MitoTracker Red. Scale bar, 10 μm. (B) The abnormal rate of mitochondrial distribution was recorded in aged control and CNP treatment oocytes. **, significant difference (P < 0.01) (C) ATP levels were measured in aged control and CNP treatment oocytes. *, significant difference (P < 0.05). (D) Mitochondrial membrane potential (ΔΨm) was detected by JC-1 staining in control and CNP treatment oocytes. Scale bar, 20 μm. (E) The ratio of red to green fluorescence intensity was calculated in aged control and CNP treatment oocytes. ****, significant difference (P < 0.0001).

**Figure S6**

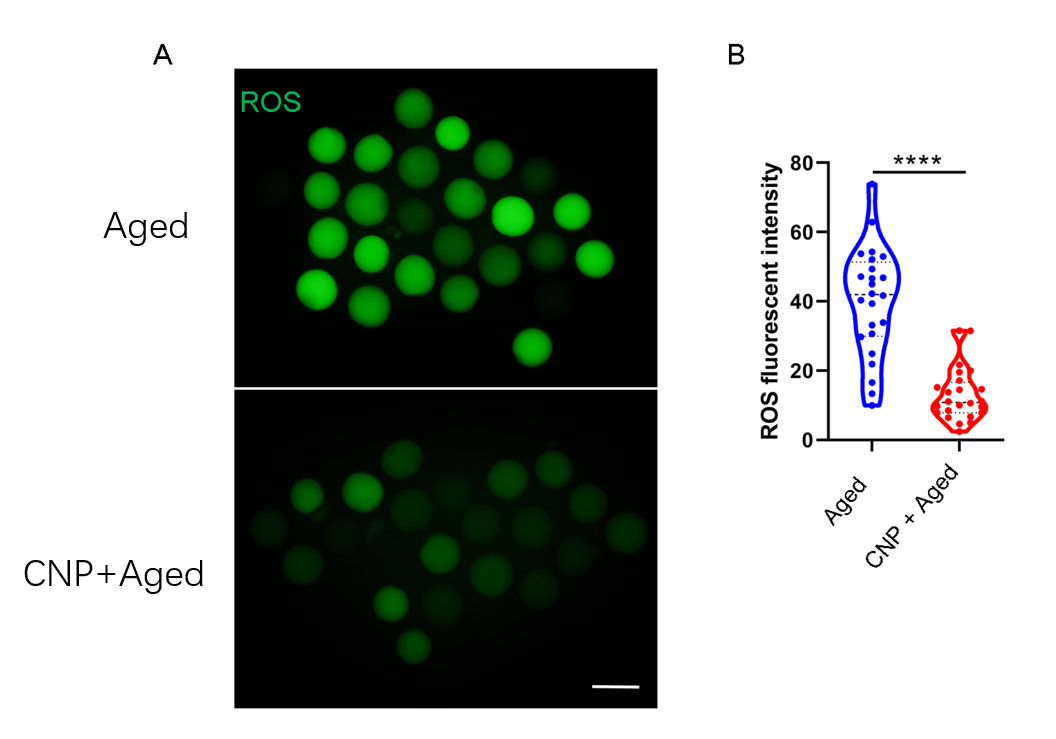

**Figure S6. Effects of CNP on the ROS content in aged oocytes.** (A) Representative images of ROS levels detected by DCFH staining in the aged control and CNP treatment oocytes. Scale bar, 100 μm. (B) The fluorescence intensity of ROS signals was measured in the aged control and CNP treatment oocytes. ****, significant difference (P < 0.0001).

**Figure S7**

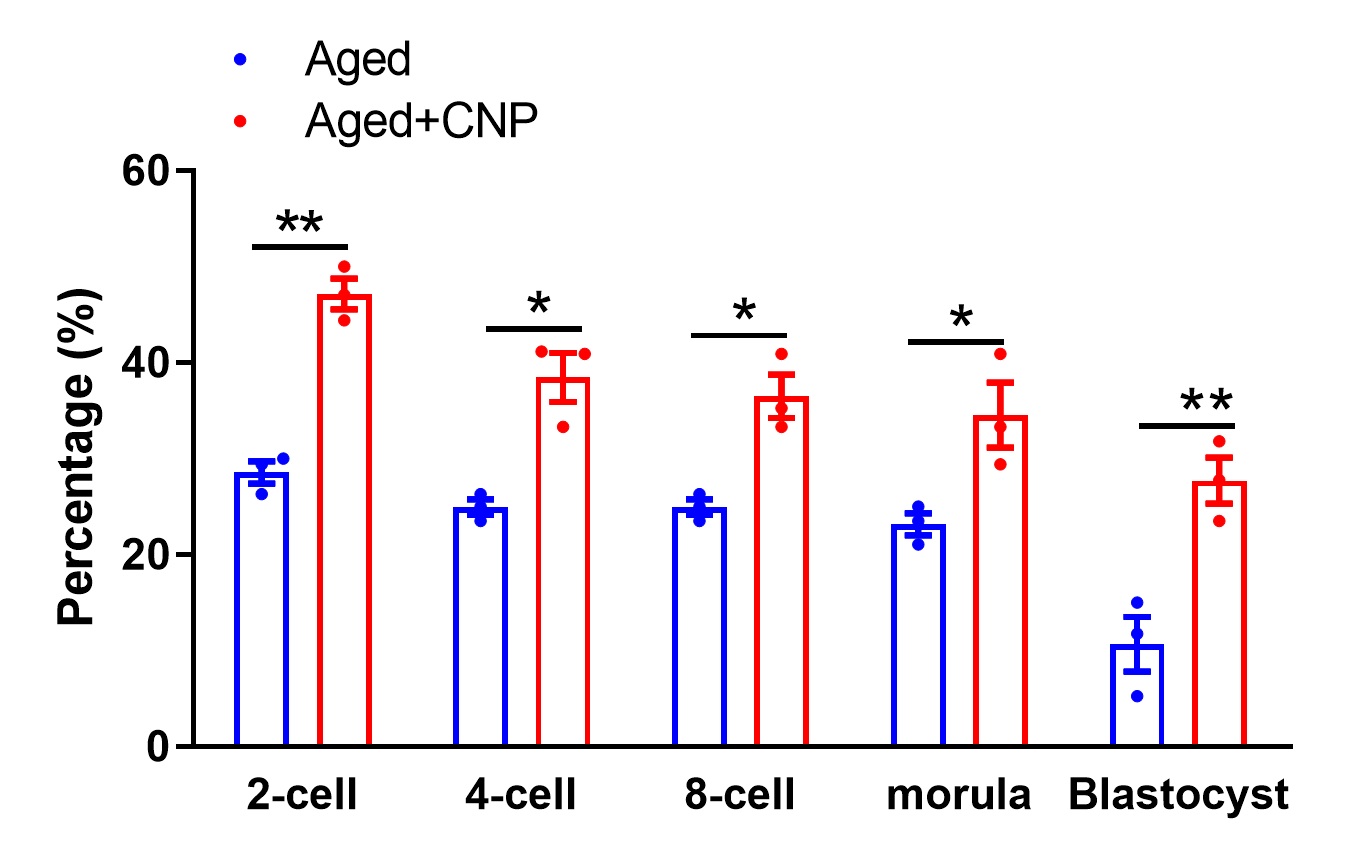

**Figure S7**. **Effects of CNP on the fertilization ability and** **embryonic development in vitro maturation oocytes.**  Data are presented as mean percentage (mean ± SEM) of three independent experiments. *p < 0.05, **p < 0.01.

**Figure S8**

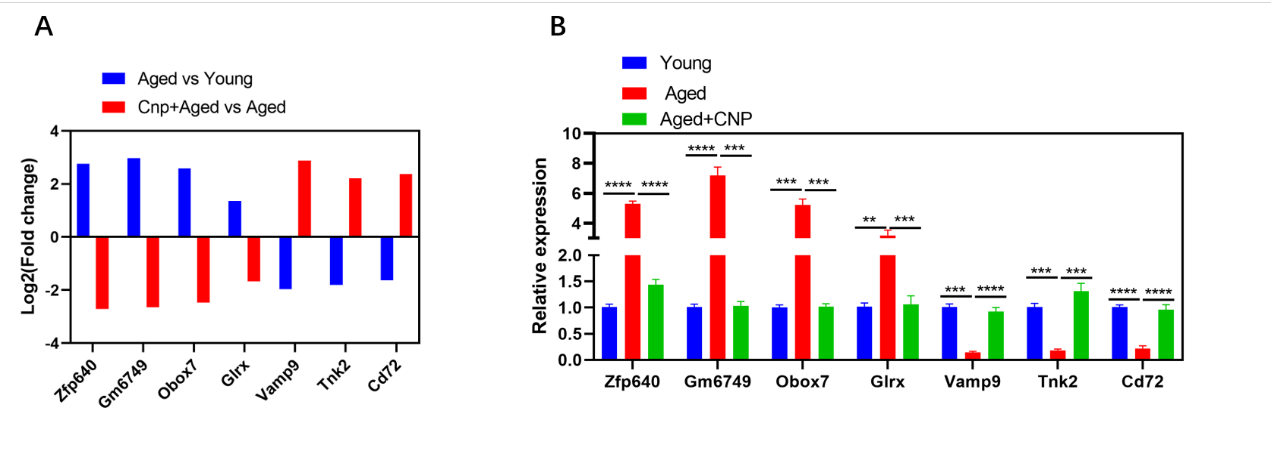

**Figure S8. The relative expression of several randomly selected genes from each group was verified using quantitative real-time PCR.**

(A) RNA-seq results of selected genes in oocyte from young, aged and aged+CNP mouse. (B) The relative expression of the randomly selected genes was verified by RT-qPCR in oocyte from young, aged and aged+CNP mouse.
